# International lineages of *Salmonella enterica* serovars isolated from chicken farms, Wakiso District, Uganda

**DOI:** 10.1101/707372

**Authors:** Takiyah A. Ball, Daniel F. Monte, Awa Aidara-Kane, Jorge Matheu, Hongyu Ru, Siddhartha Thakur, Francis Ejobi, Paula J. Fedorka-Cray

## Abstract

The growing occurrence of multidrug-resistant (MDR) *Salmonella enterica* in poultry has been reported with public health concern worldwide. We reported, recently, the occurrence of *Escherichia coli* and *Salmonella enterica* serovars carrying clinically relevant resistance genes in dairy cattle farms in the Wakiso District, Uganda, highlighting an urgent need to monitor food-producing animal environments. Here, we present the prevalence, antimicrobial resistance, and sequence type of 51 *Salmonella* isolates recovered from 400 environmental samples from chicken farms in Uganda. Among the *Salmonella* isolates, 32/51 (62.7%) were resistant to at least one antimicrobial, and 10/51 (19.6%) displayed multiple drug resistance. Through PCR, five replicon plasmids were identified among all chicken *Salmonella* including *Inc*FIIS 17/51 (33.3%), *Inc*I1α 12/51 (23.5%), *Inc*P 8/51 (15.7%), *Inc*X1 8/51 (15.7%), and *Inc*X2 1/51 (2.0%). In addition, we identified replicons through WGS (ColpVC and *Inc*FIB). A significant seasonal difference between chicken sampling periods was observed (p= 0.0017). We conclude that MDR *Salmonella* highlights the risks posed to the animals, environment, and humans for infection. Implementing a robust integrated surveillance system in Uganda will help monitor MDR to help control infectious threats.

## Introduction

Multidrug-resistant (MDR) *Salmonella enterica* remains a major public health concern being reported in food, animal, human and environmental settings, particularly in developing countries. Additionally, international lineages have been readily spread worldwide (1–6), leading a high impact on public health, which has been deemed a global pressure (WHO).

In Uganda, antibiotics are increasingly being used and not monitored or regulated in food-producing animals. This practice is well established to select antibiotic-resistant strains that can spread to humans through the food chain. In this concern, considering the lack of information regarding antimicrobial resistance (AMR) in developing countries, Uganda has plans for an integrated national surveillance system for foodborne pathogen which is included in the National Action Plan (NAP) on AMR, using a One Health approach (7).

Therefore, we present a cross-sectional study developed in chicken farms, in Uganda to investigate the prevalence, AMR, and their genomic aspects of *Salmonella enterica* serovars.

## Methods

### Bacterial Isolates

In our previous study, we reported on the phenotypic characterization of *Salmonella* isolates from cattle farms. We also collected *Salmonella* isolates from chicken farms in parallel to the cattle farms (5). This study was designed as two cross-sectional studies over one year. Sampling occurred over two seasons, the rainy season that began in March ending in September, and the dry season that began in June ending in December. Enrollment in the study occurred through contact with producers throughout the Wakiso district. A total of 20 chicken producers (20 farms) agreed to participate in the study. On-farm sampling was conducted once during the rainy and dry seasons totaling 39 collection periods (two farms dropped out of study in the rainy season). Ten samples per farm were collected at each visit totaling 379 samples (one farm had nine samples).

Ten drag swabs were used per farm. Drag swabs (3” × 3” sterile gauze pads) in sterile skim milk was the preferred collection tool (Hardy Diagnostics, Inc., Santa Maria, CA). A sampling schematic was pre-drawn to ensure maximum sampling of the house floor environment, including inside diagonals, feeding and water containers, coops, and wall to wall samples. Swabs were individually placed in a sterile whirl-pak bag; the bag was kept on ice in a cooler prior to transport to the laboratory. Isolation of *Salmonella* was collected as previously described in Fedorka-Cray et al. (8).

### Antimicrobial Resistance testing

A total of 51 *Salmonella* were isolated from chicken farms and tested for AMR using the National Antimicrobial Resistance Monitoring System (NARMS) gram-negative panels (Thermo Fisher Scientific Inc, Waltham, MA) as described by Ball et al. (5). All 51 isolates were frozen in LB broth with 30% glycerol (Thermo Fisher Scientific Inc, Waltham, MA) at −80^°^C.

### Molecular characterization

The 51 *Salmonella* isolates were struck for isolation from the frozen stocks to Tryptic Soy Agar (TSA) with 5% sheep blood (BAP) (Thermo Fisher Scientific Inc, Waltham, MA) and incubated overnight at 37^°^C to ensure purity. Lysates were prepared by suspending a loopful of well-isolated colonies into 200 μl of molecular grade water and vortexed at maximum speed for several seconds. The suspension was boiled at 100°C for 10 minutes, centrifuged at 13 × 1000 rpm for 60 seconds, and the supernatant was collected for use as the DNA template. For PCR screening and whole genome sequencing, all methods were followed as described in Ball et al. (manuscript submitted).

### Whole-genome sequencing

DNA extraction was performed using a commercial kit (QiAmp tissue, Qiagen, Germany) according to manufacturer’s guidelines. Genomic DNA (*n*= 51) were sequenced at a 300-bp paired-end-read using the Nextera XT library preparation kit at the MiSeq platform (Illumina, San Diego, CA). *De novo* assembly was achieved using CLC Genomics Workbench 10.1.1 (Qiagen). Resistome, plasmidome and multilocus sequence typing (MLST) were identified using multiple databases as ResFinder 3.1, PlasmidFinder 2.0, and MLST 2.0, respectively, available from the Center for Genomic Epidemiology (http://genomicepidemiology.org/). Sequence data were deposited in the GenomeTrakr Project.

### Statistical Analysis

The prevalence of *Salmonella* were analyzed using WHONET and Microsoft Excel. A logistic regression model was used in SAS^®^software (SAS^®^Cary, NC) where season (rainy and dry) served as the factor. Farm was included as a random effect.

## Results

Table 1 displays the results of serotype, AMR phenotype, AMR genotype, and plasmid identification. Fifty-one *Salmonella* were isolated (51/379; 13.5%) from chicken belonging eight different serotypes in order of highest to lowest, *Salmonella* serovar Enteritidis (31.3%); *S*. Kentucky (21.6%); *S*. Zanzibar and *S.* Virchow (15.7%); *S.* Newport and *S.* serovar 42:r:- (5.88%), *S*. Typhimurium (4%) and *S*. Barranquilla at 2.0%. The prevalence of *Salmonella* was statistically significantly higher in the rainy season (p=0.0017).

**Table 1:** Antimicrobial resistance phenotype and genotype comparison of Salmonella from chickens in the Wakiso district of Uganda (n=51)

| Sample ID | Serovar | ST | Resistance profile (MIC) | Resistance genes | *gyrA* | *parC* | Plasmids | Plasmids (PCR) |
| --- | --- | --- | --- | --- | --- | --- | --- | --- |
| SALM-44 | 42:r:- | 1208 | STR | Pansusceptible | none | none | Col440I | none |
| SALM-46 | 42:r:- | 1208 | STR | Pansusceptible | none | none | none | none |
| SALM-51 | 42:r:- | 1208 | STR | Pansusceptible | none | none | none | none |
| SALM-47 | Barranquilla | 3807 | Pansusceptible | Pansusceptible | none | none | none | none |
| SALM-1 | Enteritidis | 11 | Pansusceptible | *strA, strB, aadA1, blaTEM-1B, sul2, sul3, tetA* | none | none | IncFII(S), IncFIB (S), ColpVC | IncFII(S) |
| SALM-2 | Enteritidis | 11 | Pansusceptible | Pansusceptible | none | none | IncFII(S), IncFIB (S), ColpVC | IncFII(S) |
| SALM-3 | Enteritidis | 11 | Pansusceptible | *sul2* | none | none | IncFII(S), IncFIB (S), ColpVC | IncFII(S) |
| SALM-5 | Enteritidis | 11 | Pansusceptible | Pansusceptible | none | none | IncFII(S), IncFIB (S), ColpVC | IncFII(S) |
| SALM-7 | Enteritidis | 11 | Pansusceptible | Pansusceptible | none | none | IncFII(S), IncFIB (S), ColpVC | IncFII(S) |
| SALM-26 | Enteritidis | 11 | Pansusceptible | Pansusceptible | none | none | IncFII(S), IncFIB (S) | IncFII(S) |
| SALM-28 | Enteritidis | 11 | Pansusceptible | Pansusceptible | none | none | IncFII(S), IncFIB (S) | IncFII(S) |
| SALM-29 | Enteritidis | 11 | Pansusceptible | Pansusceptible | none | none | IncFII(S), IncFIB (S) | IncFII(S) |
| SALM-30 | Enteritidis | 11 | Pansusceptible | Pansusceptible | none | none | IncFII(S), IncFIB (S) | IncFII(S) |
| SALM-31 | Enteritidis | 11 | Pansusceptible | Pansusceptible | none | none | IncFII(S), IncFIB (S) | IncFII(S) |
| SALM-32 | Enteritidis | 11 | Pansusceptible | Pansusceptible | none | none | IncFII(S), IncFIB (S) | IncFII(S) |
| SALM-33 | Enteritidis | 11 | Pansusceptible | Pansusceptible | none | none | IncFII(S), IncFIB (S) | IncFII(S) |
| SALM-41 | Enteritidis | 11 | Pansusceptible | Pansusceptible | none | none | IncFII(S), IncFIB (S), Col440I | none |
| SALM-42 | Enteritidis | 11 | Pansusceptible | Pansusceptible | none | none | none | IncFII(S) |
| SALM-43 | Enteritidis | 11 | Pansusceptible | Pansusceptible | none | none | IncFII(S), IncFIB (S), Col440I | IncFII(S) |
| SALM-49 | Enteritidis | 11 | Pansusceptible | Pansusceptible | none | none | IncFII(S), IncFIB (S) | none |

Table 1 cont'd
| Sample ID | Serovar | ST | Resistance profile (MIC) | Resistance genes | *gyrA* | *parC* | Plasmids | Plasmids (PCR) |
| --- | --- | --- | --- | --- | --- | --- | --- | --- |
| SALM-8 | Kentucky | 198 | AMP, CIP, NAL, STR, SOX, TCY, SXT | *aadA1*, *aph(6)-Id*, *strA*, *strB*, *blaTEM-1B*, *dfrA14*, *qacL*, *sul2*, *sul3*, *tet(A)* | S83F/D87N | S80I | ColpVC, Inc11 | Inc11α |
| SALM-9 | Kentucky | 198 | CIP, NAL | Pansusceptible | S83F/D87N | S80I | ColpVC | none |
| SALM-10 | Kentucky | 198 | CIP, NAL, STR, SOX, TCY | *aph(3'')-Ib*, *aph(6)-Id*, *strA*, *strB*, *sul2*, *tet(A)* | S83F/D87N | S80I | ColpVC | none |
| SALM-11 | Kentucky | 198 | CIP, NAL, STR, SOX, TCY | *aph(3'')-Ib*, *aph(6)-Id*, *strA*, *strB*, *sul2*, *tet(A)* | S83F/D87N | S80I | ColpVC | none |
| SALM-12 | Kentucky | 198 | CIP, NAL, STR, SOX, TCY | *aph(3'')-Ib*, *aph(6)-Id*, *strA*, *strB*, *sul2*, *tet(A)* | S83F/D87N | S80I | ColpVC | none |
| SALM-13 | Kentucky | 198 | CIP, NAL, STR, SOX, TCY | *aph(3'')-Ib*, *aph(6)-Id*, *strA*, *strB*, *sul2*, *tet(A)* | S83F/D87N | S80I | ColpVC | none |
| SALM-14 | Kentucky | 198 | CIP, NAL, STR, SOX, TCY | *aph(3'')-Ib*, *aph(6)-Id*, *strA*, *strB*, *sul2*, *tet(A)* | S83F/D87N | S80I | ColpVC | none |
| SALM-15 | Kentucky | 198 | CIP, NAL, STR, SOX, TCY | *aph(3'')-Ib*, *aph(6)-Id*, *strA*, *strB*, *sul2*, *tet(A)* | S83F/D87N | S80I | ColpVC | none |
| SALM-16 | Kentucky | 198 | CHL, CIP, NAL, STR, SOX, TCY, SXT | *qnrS1*, *aadA1*, *aadA2*, *aph(6)-Id*, *strA*, *strB*, *cmlA1*, *dfrA14*, *sul2*, *sul3*, *tet(A)* | S83F/D87N | S80I | ColpVC, Inc11 | Inc11α |
| SALM-17 | Kentucky | 198 | CIP, NAL, STR, SOX, TCY | *strA*, *strB*, *sul2*, *tetA* | none | none | ColpVC | none |
| SALM-18 | Kentucky | 198 | AMP, CIP, NAL, SOX | *aadA1*, *blaTEM-1B*, *sul3* | S83F/D87N | S80I | ColpVC, Inc11 | Inc11α |
| SALM-45 | Newport | 166 | NAL, TCY | *qnrS1*, *tetA* | none | none | IncX2 | IncX2 |
| SALM-50 | Newport | 46 | Pansusceptible | Pansusceptible | none | none | none | none |
| SALM-4 | Typhimurium | 19 | AMP, SOX | *aadA1*, *blaTEM-1B*, *qacL*, *sul3* | none | none | Inc11, IncFII(S), IncFIB (S), ColpVC | Inc11α, IncFII(S) |
| SALM-6 | Typhimurium | 19 | AMP, SOX | *aadA1*, *blaTEM-1B*, *qacL*, *sul3* | none | none | Inc11, IncFII(S), IncFIB (S), ColpVC | Inc11α, IncFII(S) |
| SALM-34 | Virchow | 16 | NAL, TCY | *tetA* | S83Y | none | none | IncP, IncX1 |
| SALM-35 | Virchow | 16 | NAL, TCY | *tetA* | S83Y | none | none | IncP, IncX1 |

Table 1 cont'd
| Sample ID | Serovar | ST | Resistance profile (MIC) | Resistance genes | *gyrA* | *parC* | Plasmids | Plasmids (PCR) |
| --- | --- | --- | --- | --- | --- | --- | --- | --- |
| SALM-36 | Virchow | 16 | NAL, TCY | *tetA* | S83Y | none | none | IncP, IncX1 |
| SALM-37 | Virchow | 16 | NAL, TCY | *tetA* | S83Y | none | none | IncP, IncX1 |
| SALM-38 | Virchow | 16 | NAL, TCY | *tetA* | S83Y | none | none | IncP, IncX1 |
| SALM-39 | Virchow | 16 | NAL, TCY | *tetA* | S83Y | none | none | IncP, IncX1 |
| SALM-40 | Virchow | 16 | NAL, TCY | *tetA* | S83Y | none | none | IncP, IncX1 |
| SALM-48 | Virchow | 16 | NAL, TCY | *tetA* | S83Y | none | none | IncP, IncX1 |
| SALM-19 | Zanzibar | 466 | TCY | *tetA* | none | none | Incl1 | Incl1α |
| SALM-20 | Zanzibar | 466 | TCY | *tetA* | none | none | Incl1 | Incl1α |
| SALM-21 | Zanzibar | 466 | TCY | *tetA* | none | none | ColpVC, Incl1 | Incl1α |
| SALM-22 | Zanzibar | 466 | TCY | *tetA* | none | none | Incl1 | Incl1α |
| SALM-23 | Zanzibar | 466 | TCY | *tetA* | none | none | ColpVC, Incl1 | Incl1α |
| SALM-24 | Zanzibar | 466 | Pansusceptible | Pansusceptible | none | none | ColpVC | Incl1α |
| SALM-25 | Zanzibar | 466 | TCY | *tetA* | none | none | Incl1 | Incl1α |
| SALM-27 | Zanzibar | 466 | TCY | *tetA* | none | none | Incl1 | Incl1α |

The AMR phenotype displayed resistance to eight antimicrobials including tetracylcine (51%), nalidixic acid (37.3%), sulfisoxazole (23.5%), ciprofloxacin (21.6%), streptomycin (13.7%), ampicillin (7.8%), sulfamethoxazole (3.9%), chloramphenicol (2%). Whole genome sequencing analysis revealed the presence of resistance genes to tetracycline [*tetA*; 53%], sulfonamides [*sul2* (21.5%); *sul3* (11.7%)], streptomycin [*strA* (19.6%); *strB* (19.6%)], aminoglycosides [*aph(6)-Id* (15.6%); *aph(3′′)-Ib* (11.7%); *aadA1* (11.7%); *aadA2* (2%)], β-lactams [*bla*_TEM-1B;_ 9.8%], quaternary ammonium [*qacL*; 5.8%], quinolones [*qnrS1*; 5.8%] and trimethoprim [*dfrA14*; 4%]. Other than acquiring resistance genes were assigned as quinolone resistance determining regions (QRDR) with point mutation in *gyrA* and *parC* as we can observe in Table 1. Ten isolates (19.6%) showed DNA gyrase (GyrA-S83F-D87N) with a double amino acid mutation in GyrA, serine to phenylalanine at codon 83 and aspartic acid to asparagine at 87, whereas eight isolates (15.6%) showed a single amino acid substitution of serine to tyrosine at codon 83. For QRDR in *parC* was observed (n=10; 19.6%) only one substitution in serine to isoleucine at codon 80. No mutations were found in *gyrB* and *parE*.

Afterward, the prevalence of plasmids related to resistance or virulence factors were screened through sequences. Six plasmids were identified being IncFII(S)-IncFIB (S)-ColpVC the most commons distributed in S. Enteritidis; Incl1-ColpVC in *S*. Kentucky and *S.* Zanzibar; IncX2 in *S*. Newport; Incl1-IncFII(S)-IncFIB (S)-ColpVC in *S.* Typhimurium and Col440I in *S.* serovar 42:r:-.

In addition, nine sequence types (ST) such as ST11, ST198, ST466, ST16, ST166, ST46, ST19, ST1208 and ST3807 were associated with *S*. Enteritidis, *S*. Kentucky, *S*. Zanzibar, *S*. Virchow, *S*. Newport, *S*. Newport, *S*. Typhimurium, *S*. serovar 42:r:- and *S*. Barranquilla, respectively.

## Discussion

The percent prevalence of *Salmonella* (13.5%) in this study highlights the potential risk to the cross-contamination between human and poultry in Ugandan households. There are limited reports on the prevalence of *Salmonella* on chicken farms and the reports that are available show very little resistance compared to this study. Afema et al. reported 6.6% *Salmonella* was detected in live birds markets within Kampala, Uganda (9). We also learned that there was a seasonal effect in the recovery of *Salmonella*. Uganda typically has a rainy season that occurs between March to May and October to December (10). For recovery from chicken farms, a significant difference (p=0.0017) for recovery of *Salmonella* between the rainy and dry seasons as a higher prevalence of *Salmonella* was observed. During the rainy season, there is an increase in humidity as well as moisture which has been reported to influence the recovery of several bacterial species in poultry (11).

The serotype distribution in this study indicated that *Salmonella* serovars Enteritidis and Kentucky were most often recovered from chicken samples. This is comparable to the most commonly seen serotypes in chickens reported in the US (12). Kentucky has previously been reported in Uganda in humans, poultry, and the environment (9).

Among chicken isolates, *Salmonella* presented with MDR phenotypes to the antimicrobials tested. Approximately 38% of the isolates were resistant to two or more classes of antimicrobials, including two isolates resistant to seven antimicrobials. The *Salmonella* serovar Kentucky isolates in this study presented MDR to over five (ciprofloxacin, nalidixic acid, streptomycin, sulfisoxazole, and tetracycline) or seven (chloramphenicol, ampicillin, ciprofloxacin, nalidixic acid, streptomycin, sulfisoxazole, tetracycline, and trimethoprim-sulfamethoxazole) antimicrobials. All *Salmonella* serovar Kentucky isolates resistant to ciprofloxacin. Since the early 2000s, ciprofloxacin resistance for *Salmonella* serovar Kentucky has been on the rise, especially from travelers to northern and eastern Africa (13). Rickert-Hartman et al. found that 9% of the *Salmonella* serovar Kentucky isolated from travelers were ciprofloxacin resistant. An interesting note was that poultry was thought to be a reservoir for these resistant strains (13, 14). Cases of ciprofloxacin-resistant Kentucky have been seen in the US from travelers from India, resulting in seven infected with one death (13). In this regard, the emergence of *S*. Kentucky ST198 pose a major threat to public health worldwide, particularly for being highly drug-resistant (15) and has been reported in different sources including retail chicken carcasses (16). Additionally, the presence of chromosome mutation can be useful for tracking the pandemic ciprofloxacin-resistant S. Kentucky strain ST198 from geographically distinct regions (15).

We further characterized these isolates with WGS to see if concordance was seen and if isolates presented β-lactamase resistance genes. TEM-1B was identified in five isolates that PCR methods did not identify. In previous studies (17), discrepancies were seen between phenotypic resistance and genotypic analysis using WGS. It was reported that a MIC might not reach the breakpoint, but resistance genes were present (17).

Five of the 28 plasmids that were screened through PCR were observed in multiple isolates: *Inc*FIIS (17/51; 33.3%), *Inc*I1α (12/51; 23.5%), *Inc*P (8/51; 15.7%), *Inc*X1 (8/51; 15.7%), and *Inc*X2 (1/51; 2.0%). After analyzing the WGS sequences for plasmids, we notice a difference in the plasmids that were identified. In 12 isolates, there was concordance with the IncI1α, with seven of the 12 having an additional plasmid (ColpVC) that was not screened in the PCR and two with *Inc*FIIS plasmid. Seventeen isolates were in concordance with the *Inc*FIIS plasmid. These same 17 isolates also presented *Inc*FIB (S) plasmids, and ColpVC and Col4401 were identified in seven and two isolates, respectively. *Inc*X2 and *Inc*P were not identified in the WGS analysis as was in the PCR. Ten isolates were negative for PCR, but WGS identified as ColpVC (nine isolates) and Col4401 (one isolate).

*Inc*FIIS was the most common plasmid identified in this study at 33.3% (17/51) Studies have shown that bacterial isolates containing *bla*_CTX-M-1_, harbor the *Inc*FIIS along with other incompatibility plasmids (18). *Inc*1 plasmids are known to be distributed throughout many serotypes of *Salmonella* and predominate in both *E.coli* and *Salmonella* (19-21). In this study, *Inc*1α was observed among *Salmonella* serovars such as Zanzibar, Kentucky, and Typhimurium. All isolates from *Salmonella* serovar Kentucky came from the same farm, as well as isolates with *Salmonella* serovar Typhimurium.

*Inc*P and *Inc*X1 were the next most common plasmids seen in this study through PCR. Both were present in the *Salmonella* serovar Virchow isolates. It has been reported that *Inc*P can spread through groups of bacteria via conjugative transfer and code for broad range antimicrobial resistance. *Inc*P is highly likely to be found in manure, wastewater, and soil (22). *Inc*X1 is commonly found as a narrow host-range plasmid in *Enterobacteriaceae,* also spreading to other bacteria via conjugative transfer (23).

## Conclusion

In summary, we present in this study the clonal distribution of eight *Salmonella* enterica serovars displaying resistance to clinically important antibiotics. Of these, the presence of international lineages as ciprofloxacin-resistant *S*. Kentucky sequence type 198 in chicken farms raises a public concern; given that fluoroquinolones are the first treatment choice. Our findings suggest that endemic dissemination of resistant serovars, adding valuable information in the epidemiological surveillance in Uganda. Therefore, these results may encourage addition genomic surveillance studies in this region to aid the development of mitigation strategies to limit the global distribution of these multi-drug resistant *Salmonella enterica*.

## Acknowledgments

We would like to acknowledge our funding sources from North Carolina State University (NCSU), College of Veterinary Medicine and the WHO AGISAR Secretariat; Our colleagues at Makerere University Dr. Eddie Wampande, Sarah Tegule, Samuel Maling, David Apollo Munanura, Allan Odeke, Disan Muhangazi, Mark Ogal, Mutumba Paul, and Elizabeth Basemera; Our colleagues at the NCSU, College of Veterinary Medicine Diagnostic Laboratory, Dr. Megan Jacob.

